# Negative feedback control of hunger circuits by the taste of food

**DOI:** 10.1101/2023.11.30.569492

**Authors:** Tara J. Aitken, Truong Ly, Sarah Shehata, Nilla Sivakumar, Naymalis La Santa Medina, Lindsay A. Gray, Naz Dundar, Chris Barnes, Zachary A. Knight

## Abstract

The rewarding taste of food is critical for motivating animals to eat, but whether taste has a parallel function in promoting meal termination is not well understood. Here we show that hunger-promoting AgRP neurons are rapidly inhibited during each bout of ingestion by a signal linked to the taste of food. Blocking these transient dips in activity via closed-loop optogenetic stimulation increases food intake by selectively delaying the onset of satiety. We show that upstream leptin receptor-expressing neurons in the dorsomedial hypothalamus (DMH^LepR^) are tuned to respond to sweet or fatty tastes and exhibit time-locked activation during feeding that is the mirror image of downstream AgRP cells. These findings reveal an unexpected role for taste in the negative feedback control of ingestion. They also reveal a mechanism by which AgRP neurons, which are the primary cells that drive hunger, are able to influence the moment-by-moment dynamics of food consumption.

## Introduction

All animals face the basic challenge of regulating the size of each meal.^1^ This regulation is thought to be mediated by the balance between two opposing forces: the sense of taste, which provides the positive feedback that propels the meal forward (i.e. we eat because food tastes good), and visceral feedback from the stomach and intestines, which provides the negative feedback that drives the termination of feeding.

Nevertheless, the fact that nutrient sensing in the intestine is inherently limited by slow gastric emptying^2^ raises the question of whether other chemosensory signals also contribute to satiation, possibly by providing an early estimate of food consumed. Several observations suggest that taste cues could play this role, albeit through mechanisms that are not well understood. One line of evidence comes from studies showing that food is more satiating when consumed by mouth than when delivered directly to the stomach or intestines.^3–7^ Indeed, humans receiving enteral tube feeding report a need to chew their food in order to fully quell their hunger, even if the tasted food is never actually swallowed.^5,8^ A second line of evidence comes from studies that used sham feeding in rats, a preparation in which ingested food is allowed to drain out of the stomach, thereby eliminating all GI feedback.^9^ This revealed that an important component of satiation is linked to the taste of food and remains intact even when all GI signals are lost.^10–12^ Finally, sensory-specific satiety is the phenomenon whereby repeated exposure to the same taste can result in early termination of its consumption, independent of any post-ingestive feedback.^13,14^ For all of these phenomena, it is thought that appetitive food tastes – sweet and fat – act as a negative feedback signal that contributes to the termination of a normal meal, and that they do so by a mechanism independent of any innate or conditioned aversion.

The neural mechanisms that underlie these phenomena are completely unknown. A fundamental challenge is that any manipulation of taste itself will impair not only these negative feedback mechanisms that contribute to satiation, but also the positive feedback (i.e. food reward) that is essential for the initiation and maintenance of ingestion in the first place. Thus, behavioral analysis alone cannot disentangle how gustatory cues are used, simultaneously, by different brain systems for opposing purposes. Instead, unraveling this regulation will likely require identification and characterization of the specific circuits where taste signals act in the brain to produce these opposing effects on behavior.

Agouti-related peptide (AgRP) neurons in the arcuate nucleus (ARC) of the hypothalamus are a key cell type for the control of feeding behavior,^15^ and therefore a candidate site in the brain where taste cues may act to modulate ingestion. AgRP neurons are activated by food deprivation,^16,17^ and their artificial stimulation broadly recapitulates the motivational and sensory hallmarks of hunger, including avid food seeking and consumption in fed animals.^18–22^ In contrast, silencing of AgRP neurons in fasted mice attenuates many of these responses.^19,23^ Thus, AgRP neurons are pivotal for controlling the desire to eat, and investigation of their natural regulation can reveal mechanisms that control feeding behavior.

Whether and how AgRP neurons are modulated by taste and other orosensory cues is unknown, in part due to the unusual regulation of these cells. Although AgRP neurons are gradually activated during food deprivation, they are inhibited within seconds when a hungry mouse sees and smells food.^17,20,24^ This rapid, global decrease in AgRP neuron activity occurs before the first bite of food is consumed, is sustained for the duration of the meal, and anticipates the number of calories subsequently consumed.^25^ Because AgRP neuron activity is greatly reduced before ingestion begins, much less is known about the moment-by-moment dynamics of AgRP neurons during feeding itself. Similarly, the direct GABAergic inputs to AgRP neurons (DMH^LepR^ neurons) are activated by the sight and smell of food before the onset of ingestion,^26^ suggesting that the broader hunger circuit is regulated primarily in anticipation of future food consumption. These observations raise the question of whether the activity of these neurons during ingestion itself has any relevance to behavior.^21,27^

In the experiments that follow, we show that AgRP neurons receive time-locked inhibition during ingestion by a signal linked to the taste of food. Furthermore, their direct afferents, DMH^LepR^ neurons, encode a representation of food-associated tastes. Selectively blocking this gustatory feedback using closed-loop stimulation delays meal termination. These findings reveal a mechanism by which the taste of food, acting through inhibition of AgRP neurons, functions as a negative feedback signal to promote satiation.

## Results

### AgRP neurons are inhibited in a manner time-locked to ingestion

To investigate a potential role for orosensory cues inhibiting appetite, we targeted GCaMP8s to AgRP neurons and recorded their dynamics during the consummatory phase of feeding by fiber photometry (Figure 1A). Mice were fasted overnight and then given access to a sipper containing the liquid diet Ensure for self-paced feeding (Figure 1B). As expected, presentation of the sipper resulted in an immediate and sustained decrease in AgRP neuron activity (-6.0 ± 0.8 z, p = 0.0018; Figure 1B,C). However, as feeding began, we noticed an additional, time-locked inhibition of AgRP neurons that occurred during each bout of consumption (-2.5 ± 0.2 z, p = 0.0004; Figure 1B,D). This time-locked inhibition began immediately following the first lick in each bout (Figure 1C) and reached a minimum closely following the last lick in each bout (time to min 9.9 ±1.2 s; Figure 1E) before gradually returning to baseline (time constant 9.5 ±1.9 s Figure 1E). This inhibition was correlated with licking, and this correlation was abolished when the licking data was shuffled (correlation coefficient -0.206 ± 0.019 vs. 0.000122 ± 0.00261, p = 0.0004; Figure 1L). The magnitude of the lick-triggered inhibition was similar in fasted and fed animals (-3.1 ± 0.2 z fasted; -5.5 ± 0.6 z fed, p=0.43), indicating that it does not depend on nutritional state (Figure 1C-E), and was consistent throughout the trial (-5.8 ± 0.7 z first 10 minutes; -4.3 ± 0.5 z last 10 minutes, p=0.76), indicating that it does not attenuate with ingestion (Figure 1F). Thus, AgRP neurons are inhibited by a transient cue during each bout of ingestion that is triggered by licking and distinct from the well-characterized tonic inhibition that occurs at the beginning of the meal.

**Figure 1.**
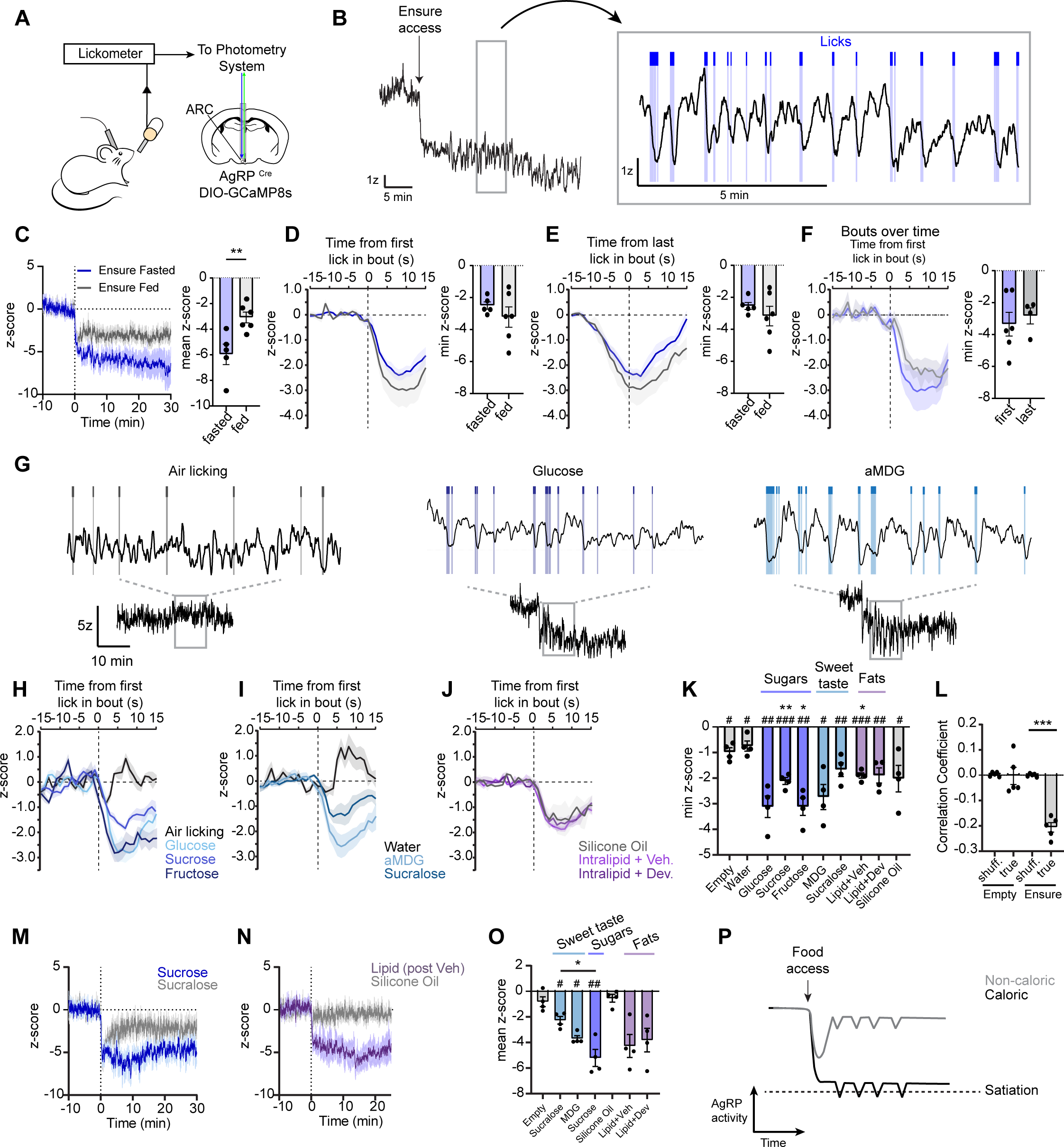
AgRP neurons track the taste of food. **A,** Schematic of mice equipped for fiber photometry of AgRP neurons while licking a bottle attached to a lickometer. **B,** Example trace of AgRP activity aligned to licks. **C,** Z-scored AgRP activity in fasted or fed mice drinking ensure. **D,** PSTH of AgRP activity and minimum z-score peri the first lick. **E,** Same as d but peri the last lick. **F,** PSTH of AgRP activity the first or last 10 minutes of drinking ensure. **G,** Example whole-trial and zoomed-in traces of AgRP activity during licking. **H-J,** PSTH of AgRP activity peri the first lick. **K,** Minimum z-score during licking bouts of the solutions in h-j. **L,** Correlation coefficient between AgRP activity and true or shuffled lick data. **M,N,** Z-scored AgRP activity during consumption of sweet (m), or fatty (n) solutions. **O,** Mean z-score over ten minutes of consumption. **P,** Proposed model of AgRP activity integrating different levels of signals to achieve satiation. MDG = alpha-methyl-d-glucopyranoside. N.S. p>0.05, *p<0.05, **p<0.01 for direct comparisons; compared to water (k). #p<0.05, ##p<0.01, ###p<0.001 relative to 0 (k). #p<0.05 relative to empty (o). Dots represent individual mice. Data is presented as mean ± SEM. See also Figure S1.

To clarify the determinants of this lick-evoked inhibition, we tested a panel of solutions that differ in their gustatory and nutritional properties. We observed no response to licking an empty bottle (Figure 1G,H) and weak activation following ingestion of water (Figure 1I), indicating that ingestion itself is insufficient for AgRP neuron inhibition. On the other hand, consumption of pure fat (Intralipid) or pure sugar (glucose) resulted in robust, time-locked inhibition of AgRP neurons during licking (-1.9 ± 0.1 z, p=0.0004, and -3.1 ± 0.4 z p=0.005 compared to H_0_=0), indicating that the response is triggered by food but is not dependent on a specific macronutrient (Figure 1K). The inhibition by sugar and fat was too fast to be mediated by post-ingestive feedback, and, consistently, it did not depend on known mechanisms by which GI nutrients are sensed and communicated to AgRP neurons.^25,28^ For example, pre-treatment with the cholecystokinin A receptor (CCKAR) antagonist devazepide, which abolishes the inhibition of AgRP neurons when Intralipid infused in the stomach,^25^ had no effect on the rapid inhibition of AgRP neurons during licking of Intralipid (Figure 1J,K). Similarly, sugars that are detected by different post-ingestive mechanisms (e.g. glucose and fructose Figure 1H,K) caused similar lick-triggered inhibition of AgRP neurons, suggesting that response is driven by their sweet taste rather than a specific gut sensor. To test the role of orosensory cues directly, we measured responses to consumption of sucralose, which is an artificial sweetener; α-methyl-D-glucopyranoside (MDG), which tastes sweet and also activates the glucose sensor SGLT1,^29,30^ but lacks calories; and silicone oil, which mimics the texture of fat but has no calories. We observed significant time-locked inhibition of AgRP neurons during ingestion of all three non-nutritive substances (Figure 1H,I,K). This inhibition was highly correlated to licking (Figure S1) and was sustained throughout the trial (Figure S1), indicating that it does not reflect, for example, momentary uncertainty about whether the ingested solutions are nutritive. Rather, this indicates that a signal tightly linked to the orosensory properties of food, but not necessarily their calorie content, transiently inhibits AgRP neurons during each bout of ingestion.

The fact that AgRP neurons are inhibited during licking of non-caloric substances is counterintuitive because these substances are not satiating. However, it is important to emphasize that the rapid inhibition of AgRP neurons that occurs at the beginning of the meal (i.e. in response to the “sight and smell” of food) is sustained only when followed by ingestion of calories^24^ and, consistently, we confirmed that the reduction in AgRP neuron activity across the entire trial (i.e. the baseline change) was much larger following consumption of nutritive foods (e.g. sucrose) than their non-nutritive counterparts (e.g. sucralose, -5.2 ± 0.7 z vs -2.3 ± 0.3 z, p = 0.047; Figure 1M-O). The one exception to this observation was MDG (mean -3.7 ± 0.22 z, p=0.047 vs empty bottle), which is non-caloric but nevertheless mimics nutrients by activating SGLT1. An important consequence of this baseline difference is that AgRP neuron activity is even further reduced when licking nutritive substances compared to their non-nutritive counterparts, even though the magnitude of the lick-triggered inhibition caused by these substances may be similar (Figure 1P). This provides a mechanism by which a signal linked purely to the taste of food could nevertheless modulate AgRP neurons, and therefore food intake, in a calorie-dependent way.

### Ingestion-triggered dips in AgRP neuron activity control meal duration

The fact that the phasic dips in AgRP neuron activity are precisely timed to bouts of ingestion suggests that they may be involved in regulating consumption. If so, this would provide a mechanism by which AgRP neuron dynamics during feeding influence ongoing behavior, which to date has been elusive. To test this, we used optogenetics to selectively block these transient dips in activity during licking and measure the effect on food intake.

Mice expressing channelrhodopsin in AgRP neurons were equipped with an optical fiber above the ARC and then given access to a lickometer containing Ensure for self-paced feeding (Figure 2A). We first confirmed that high frequency stimulation (20 Hz) of AgRP neurons in manner independent of licking (i.e. tonic) caused a dramatic increase in Ensure consumption (3518 ± 364 vs 736 ± 176 licks, with and without laser, p=0.0039). We then tested a closed-loop protocol designed to selectively reverse the ingestion-induced dips in AgRP neuron activity, but not artificially stimulate the cells to the level found in fasted animals. To do this, we programmed the laser to deliver low-frequency stimulation (5 Hz) triggered by licking and confined to ongoing lick bouts. This frequency was chosen, in part, based on data from electrophysiologic recordings showing that AgRP neurons in fasted mice have much higher tonic firing rates (in the range of 20-30 Hz).^17^ Finally, we partitioned the experiment into a series of two-minute blocks, in which blocks containing closed-loop stimulation were randomly interleaved with blocks containing no stimulation (Figure 2B). This was done to measure the duration of any behavioral effects of AgRP neuron stimulation, which is important because under certain conditions AgRP neuron stimulation can cause long-lasting increases in feeding.^21,31,32^

**Figure 2.**
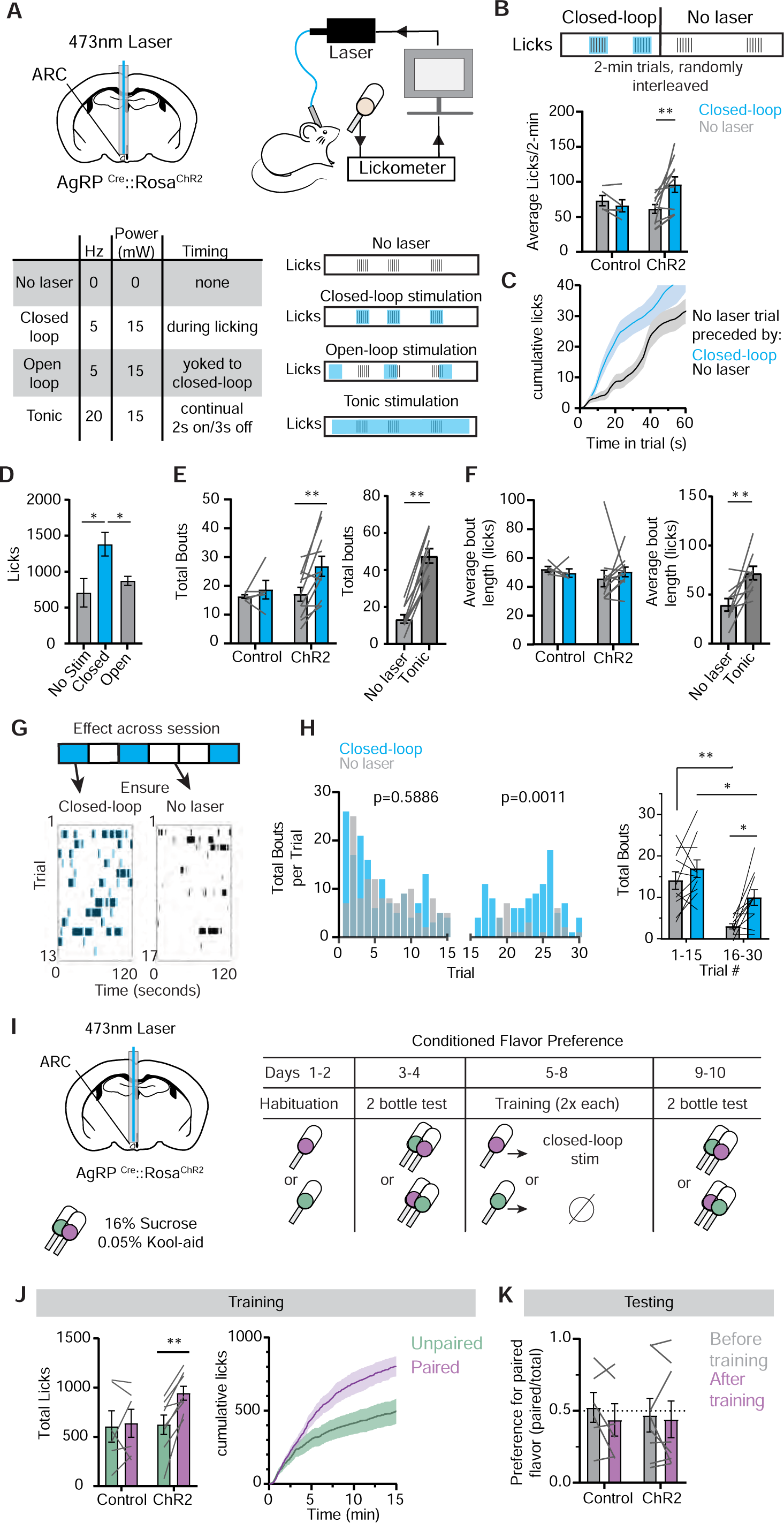
Ingestion-triggered dips in AgRP activity control meal duration. **A,** Schematic of mice equipped with an optic ferrule above the arcuate nucleus (ARC), where AgRP neurons endogenously express ChR2. Table and schematic of stimulation protocols used. **B,** Schematic of experimental design involved randomly interleaving no laser trials with closed-loop trials. Average licks per 2-min trial for ChR2 and control mice. **C,** CDF of no laser trials preceded by either a closed-loop or no laser trial. **D,** Total licks for three different protocols. **E,** Total bouts for the interleaved closed-loop protocol shown in B (left) and tonic stimulation. **F,** Same as E but for bout length. **G,** Example feeding behavior during a closed-loop session, collapsed by trial type. **H,** Total bouts across all mice, separated by trial type as a distribution (left). Total bouts for ChR2 mice split by trial type and time in session (right). **I,** Schematic of conditioned flavor preference protocol. **J,** Training data for paired and unpaired flavors. **K,** Preference for flavor (licks for paired flavor divided by total licks) before and after training. N.S. p>0.05, *p<0.05, **p<0.01. Dots represent individual mice. Data is presented as mean ± SEM.

We found that closed-loop stimulation of AgRP neurons during licking robustly increased food intake relative to control mice that lacked ChR2 expression but otherwise were treated identically ((F_ChR2 x Trial_; 1,14) = 7.12, p=0.018; Figure 2B). Strikingly, this effect was confined to the laser-paired blocks, as mice ate no more than controls during the interleaved blocks that received no laser stimulation (Figure 2B). Consistently, following a transition from a closed-loop to a no laser block, the lick rate rapidly declined (Figure 2C). This indicates that the behavioral response to low-frequency, closed-loop stimulation is temporally confined and therefore involves a mechanism that is distinct from the “sustained hunger” that is induced by tonic, high-frequency stimulation and has been reported previously.^21,32^

During closed-loop experiments, mice received only ∼4% of the laser pulses that are normally delivered during tonic stimulation of AgRP neuron (909 vs 21542 total pulses on average, p<0.0001). This suggests that the behavioral response to closed-loop stimulation depends on the precise timing of laser stimulation. To test this, we performed an open-loop stimulation experiment in which mice received the same number of laser pulses as the preceding closed-loop trial, but in a manner uncoupled from licking (i.e., randomly distributed throughout the session; Figure 2A). This open-loop stimulation had no effect on food intake (707 ± 198 licks without laser vs 872 ± 63 licks with open-loop, p=0.67; Figure 2D). This reveals that low-frequency AgRP stimulation can drive food intake only when it is precisely timed to block the natural dips that occur during ingestion.

We found that closed-loop stimulation selectively increased the number of licking bouts, with no effect on their size (bout number: 17 ± 2.42 vs 26.7 ± 3.50, p = 0.0039; bout length: 45.7 5.76 ± vs 50.27 ± 3.35 licks, p = 0.687; Figure 2E,F). In contrast, we found that high-frequency, tonic stimulation of AgRP neurons increased both the number of licking bouts (13.6 vs 47.7 bouts, no laser vs. laser; p=0.0039; Figure 2E) and their size (39.6 vs 72.0 licks, p=0.0078; Figure 2F). In rodents, the rate of bout initiation tracks changes in incentive value, which declines as a meal progresses,^33–37^ whereas the size of a licking bout tracks food palatability and is thought to reflect hedonic motivation or “liking”.^34,35,38–43^ Thus, these data suggest that the ingestion-triggered dips in AgRP neuron activity may be involved in the changes in incentive value that occur with satiation,^33,34^ whereas strong activation of AgRP neurons engages both of these motivational mechanisms.

We reasoned that if the ingestion-triggered dips in AgRP neuron activity are involved in satiation, then blocking these dips should increase feeding only later in the trial, when the animals begin to approach satiety. Indeed, we found that closed-loop stimulation selectively attenuated the decrease in bout number in the second half of the trial (3.0 ± 0.6 vs 9.9 ± 1.9 bouts, p=0.034), with no effect on the number of bouts in the first half (14 ± 2.1 vs 16.8 ± 2.1 bouts, p=0.75; Figure 2G,H). On the other hand, the fact that closed-loop stimulation did not affect bout size predicts that the ingestion-triggered dips are not involved in modulating food palatability.^34,35,40–43^ To test this, we performed a flavor conditioning experiment in which one flavor was paired with closed-loop stimulation during licking and the other was not (Figure 2I). We found that low frequency (5 Hz), closed-loop stimulation increased consumption of the paired flavor during training, confirming the efficacy of stimulation (943 ± 72 vs 622 ±99 licks, p=0.0018; Figure 2J). However, there was no effect on preference in a subsequent two-bottle test, indicating this protocol does not produce learned increases in palatability (Figure 2K). Taken together with the data from Figure 1, this reveals that a signal linked to the taste of food transiently inhibits AgRP neurons during each bout of ingestion, and that blocking these dips specifically delays the process of satiation that leads to meal termination.

### DMH^LepR^ neurons are activated in a manner time-locked to ingestion

The time-locked inhibition of AgRP neurons during licking suggests that they receive GABAergic input that relays this gustatory feedback. To identify the source of this signal, we examined leptin-receptor expressing neurons in the dorsomedial hypothalamus (DMH^LepR^ neurons), which are a major population of cells that provide direct, GABAergic input to AgRP neurons^26^ (Figure 3A). We prepared mice for single-cell calcium imaging of DMH^LepR^ neurons by targeting GCaMP6s to the DMH of LepR^Cre^ mice and installing a GRIN lens above the injection site (Figure 3B).

**Figure 3.**
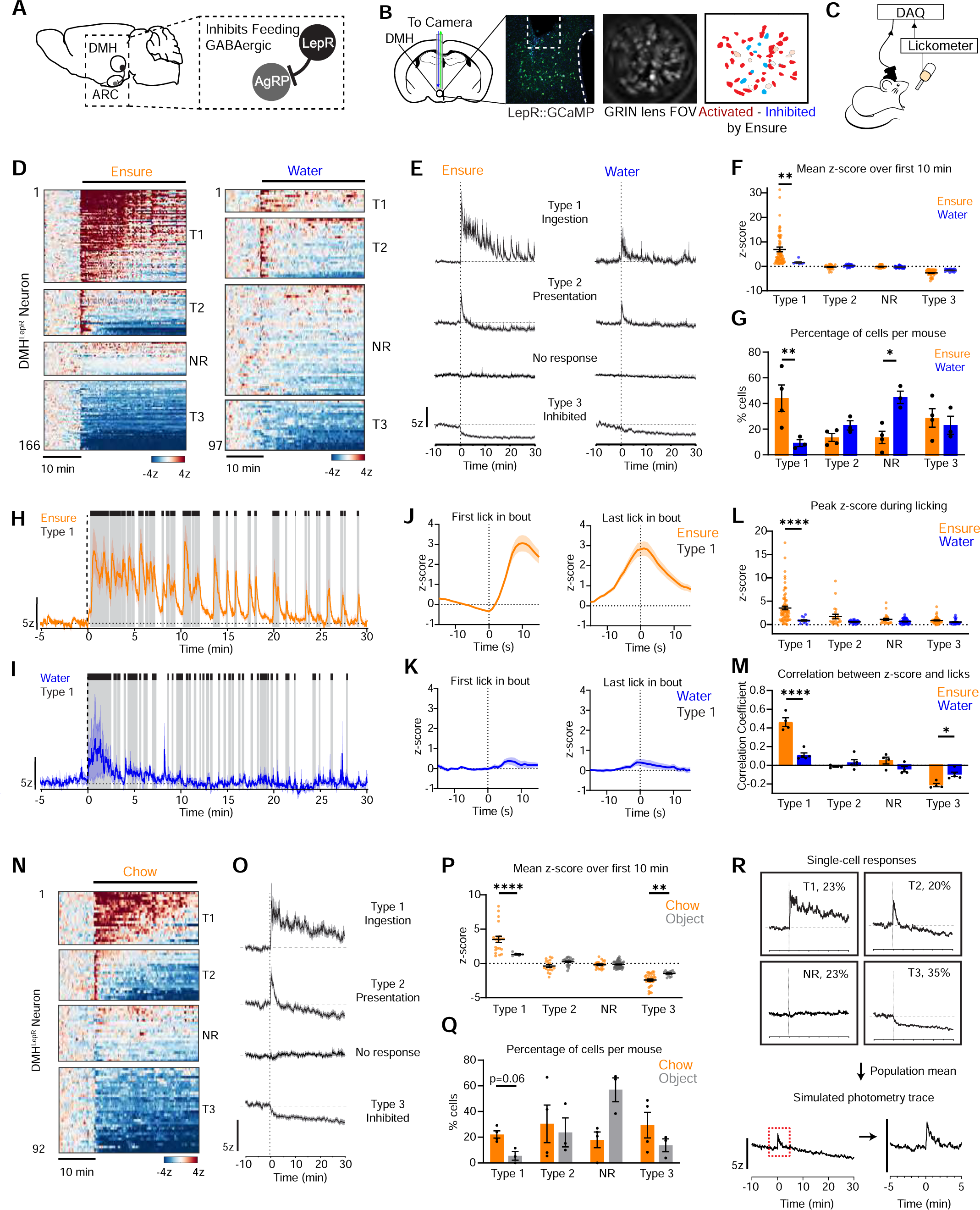
DMH^LepR^ neurons are activated time-locked to ingestion. **A**, Inhibitory circuit schematic from DMH to ARC. **B**, Schematic and example of lens placement above GCaMP-expressing DMH^LepR^ neurons. Example field of view color-coded to responses during consumption. **C**, Schematic of single-cell calcium imaging during consumption. **D**, Heatmap of DMH^LepR^ responses (N=4-5) during consumption while fasted. **E**, Averaged traces of categories in D. **F**, Mean z-score of individual neurons over first ten minutes. **G**, Percentage of each category per mouse. **H,I**, Example T1 averaged trace during licking (grey) ensure (H) or water (I). **J,K**, PSTH of T1 activity peri the first or last lick of ensure (J) or water (K). **L**, Peak z-score during licking of individual neurons. **M**, Correlation coefficient for neural activity against licks. **N**, Same as D but for chow. **O**, Same as E but for chow. **P,Q**, Same as L,M but for chow and object. **R**, Generation of pseudo-photometry trace during chow consumption. DMH = dorsomedial hypothalamus. ARC = arcuate nucleus. T1 = Type 1, T2 = Type 2, T3 = Type 3, NR = no response. N.S. p>0.05, *p<0.05, **p<0.01, ****p<0.0001. Dots represent individual mice unless otherwise noted. Data is presented as mean ± SEM. See also Figures S2 and S3.

Mice were fasted overnight and then given access to a bottle of Ensure for self-paced feeding (Figure 3C). We found that the most DMH^LepR^ neurons were strongly modulated at the start of the trial (85%, Figure 3D-G). These modulated cells fell into three categories: a major subset of activated cells (44% of all neurons) that were phasically activated throughout the trial in response to ingestion (Type 1 or “ingestion-activated” cells); a smaller subset of activated cells (13% of neurons) that were transiently activated only when the sipper was first presented (Type 2 or “presentation-activated” cells); and cells that were rapidly and durably inhibited when the trial began (28%; Type 3 or “inhibited” cells). These differentially modulated cells were anatomically intermingled at the scale of our 0.5 mm GRIN lens recordings (Figure 3B and S3).

The activity of Type 1 cells was strikingly time-locked to licking Ensure (Figure 3H-K). These cells were activated following the first lick in each bout (time to peak 10.5 ± 0.40 s) and reached a maximum at the last lick of the bout (3.3 ± 0.3 z), before gradually decaying when licking ceased (tau 8.8 ± 0.4 s). This correlation between calcium dynamics and licking was abolished when the licking data was shuffled (type 1 correlation coefficient: 0.463 ± 0.0467 vs -0.00196 ± 0.00465, p < 0.0001; Figure 3M, S2). In contrast to these robust responses to Ensure, many fewer DMH^LepR^ neurons were activated when mice drank water (44.1% vs 9.2%, p=0.0043), and the magnitude of these responses was smaller (3.6 vs 0.7 z, p<0.0001; Figure 3G-K). We also observed only minimal responses to air licking at an empty sipper (Figure S2). Thus, many LepR neurons are specifically activated by licking Ensure, with time locked dynamics that are the mirror image of downstream AgRP neurons.

Whereas the activity of Type 1 neurons was tightly linked to food consumption, both Type 2 cells (“presentation-activated”) and Type 3 cells (“inhibited”) responded similarly when mice were presented with water instead of Ensure, as measured by the percentage of modulated cells (Figure 3G) and the strength of their modulation (Figure 3F). Moreover, these cells showed minimal responses time-locked to ingestion (Figure 3L). This suggests that the activity of these other DMH^LepR^ subsets is linked to a more general signal associated with behavior, such as salience or arousal, rather than feeding per se.

In contrast to our finding that many DMH^LepR^ neurons show time-locked activation during ingestion throughout the entire meal, previous studies using fiber photometry reported that DMH^LepR^ neurons are activated only transiently when food is first presented (duration ∼1 min).^26^ We confirmed that this discrepancy was not due differences in the food stimulus, because we also observed ingestion-triggered activation of DMH^LepR^ neurons when mice consumed chow (Figure 3N-Q) or peanut butter (Figure S2). To test whether this discrepancy was due to differences in recording method, we collapsed our single-cell calcium imaging data into a mean response that mimics a photometry trace (Figure 3R). The resulting mean trace shows a transient activation upon food presentation (duration ∼2 min) that closely resembles what has previously been reported by photometry.^26,44^ This suggests that the time-locked activation of DMH^LepR^ neurons during ingestion – which is the major response of these cells to food – may have been overlooked in prior studies due to population averaging by photometry.

### DMH-LepR neurons are activated by the taste of food

We sought to identify the signals that drive the time-locked activation of DMH^LepR^ neurons during feeding. To isolate the contribution of orosensory cues, we used a Davis Rig^45^ to perform brief access taste tests, in which mice are given access to a variety of tastants for five seconds each in pseudorandom order (Figure 4A). An important advantage of this approach is that it minimizes the contribution of gastrointestinal feedback (because the total amount ingested is small) and exterosensory cues (because animals have limited ability predict the upcoming tastant based on sight or smell).

**Figure 4.**
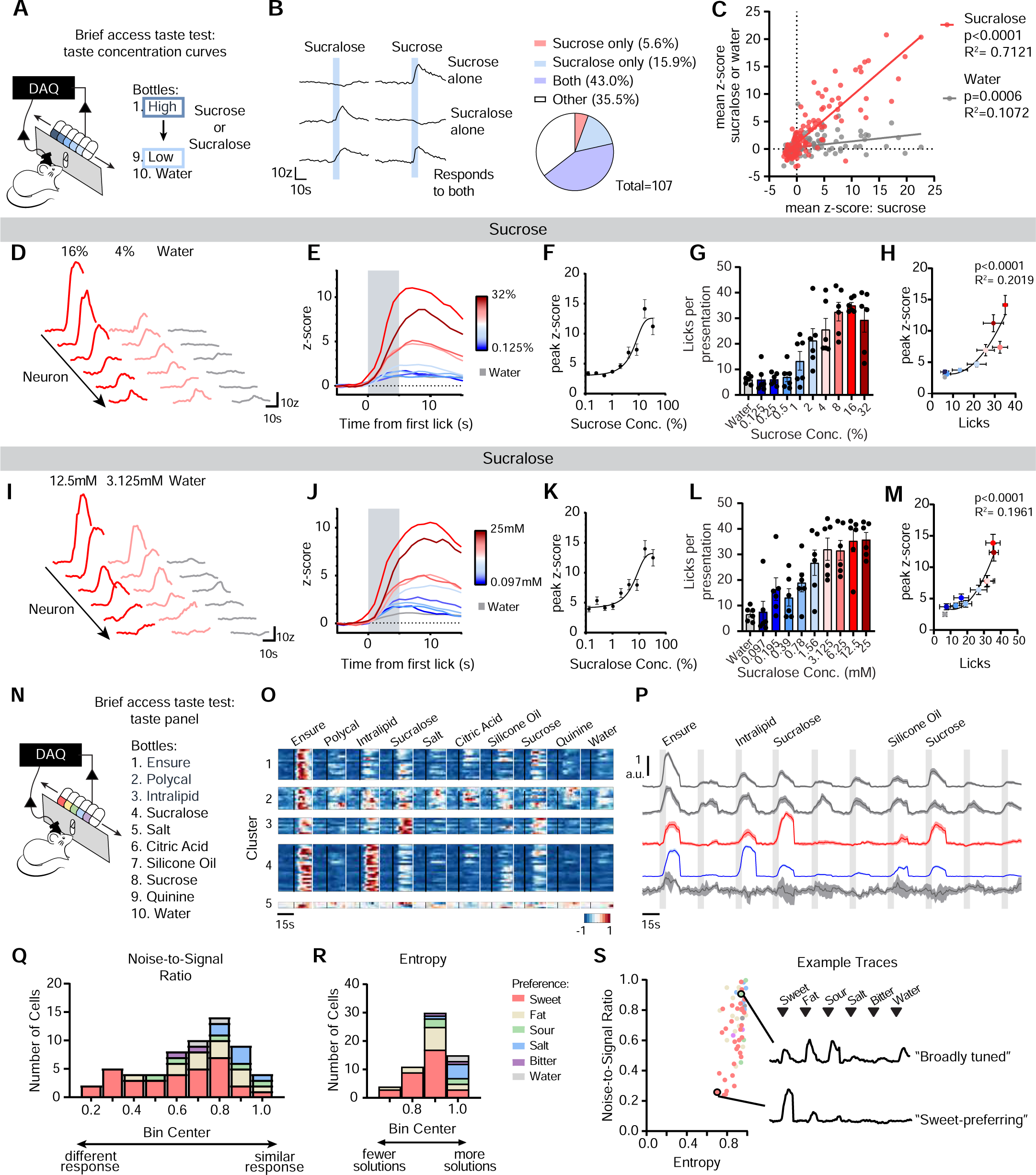
DMH^LepR^ neurons are activated by the taste of food. **A**, Diagram of Davis Rig used for brief access taste tests during single-cell imaging of DMH^LepR^ neurons. **B**, Example traces of neurons preferring sucrose, sucralose, or both, and corresponding quantification. **C**, Correlation plot of activated neurons comparing the mean z-score for sucrose against either sucralose or water. Each point represents a single neuron. P-value indicates significance relative to a slope of 0. **D,** Example responses to sucrose concentrations. **E,** Mean traces of neurons activated by second-to-maximal concentration across all concentrations of sucrose. **F,** Peak z-score across sucrose concentrations. **G,** Licks per presentation. **H**, non-linear regression between peak z-score and licks. P-value indicates significance of fit relative to a linear regression. **I-L**, Same as D-H but for sucralose. **N**, Schematic of setup for taste panel experiment. **O,** K-means clustering of ensure-activated neurons across all solutions, presented as one heatmap per cluster. **P,** Average traces for each cluster in O, aligned to solution access. **Q**, Noise-to-signal ratio **R**, entropy, and **S**, noise-to-signal ratio versus entropy plot of ensure-activated neurons, colored by their preferred non-caloric taste. Data is reported as the mean, with error bars as ±SEM.

Given that DMH^LepR^ neurons were strongly activated by Ensure, which is mostly sugar, we tested first the responses of these neurons to sweet tastes (Figure 4A-M). Mice equipped for single-cell imaging were fasted overnight and then presented with a panel of nine different concentrations of sucrose and water in pseudo-randomized sequence (Figure 4A). This revealed a striking dose-dependent response, in which the magnitude of neural activation scaled with the sweetness of the sucrose solution. Interestingly, the response peaked at 16% sucrose and declined at 32%, which mirrors the known sweetness preferences of mice (Figure 4D-F).^46^ Although the number of licks for each solution also scaled with sweetness, the alone cannot explain the increase in neural activity, because the lick responses plateaued at a lower sucrose concentration than the neural responses (Figure 4F,G). Consistently, the relationship between peak z-score and licks was better fit by a second order polynomial than a linear regression (Extra sum-of-squares F test, quadratic versus linear regression: F(1,677) = 15.97, p < 0.0001; Figure 4H). This reveals that DMH^LepR^ neurons are activated by sucrose ingestion in a manner that tracks sweetness and is separable from the amount consumed.

We next asked if this acute neural response to sucrose ingestion requires caloric value. To do this, we performed the analogous brief access taste test with nine different concentrations of the non-caloric sweetener sucralose. This revealed a remarkably similar dose-dependent response to sweetener concentration (Figure 4I-M). As with sucrose, the lick responses plateaued at a lower sucrose concentration than the neural responses, and the relationship between neural activity and licks was non-linear (Extra sum-of-squares F test, quadratic versus linear regression: F(1,647) = 17.58, p < 0.0001; Figure 4M). This confirms that sweetness has an effect on DMH^LepR^ neuron activity that is separable from the act of consumption.

We next compared the responses to sucrose and sucralose and asked if the same DMH^LepR^ neurons respond to both and, if so, do they respond in the same way. To examine this, we aligned neurons across trials of the most preferred sweetness for sucrose and sucralose, which revealed that most neurons that were activated by sucrose were also activated by sucralose, and vice-versa (Figure 4B). Moreover, there was a strong positive correlation between the magnitude of an individual neuron’s response to sucrose and its response to sucralose (R^2^=0.7121, p<0.0001). In contrast, the correlation between neural responses to sucrose and water was weaker (R^2^=0.1072, p=0.0006) and almost all cells responded more strongly to sucrose (Figure 4C). This further supports the idea that a major subpopulation of DMH^LepR^ neurons is specialized to track sweet taste.

The function of sweet taste is to signal that a food has calories. The orosensory detection of fat has a similar purpose, whereas other gustatory cues, such as sour, bitter and salt, have different roles in guiding feeding behavior. Given that DMH^LepR^ neurons are involved in regulating caloric hunger,^26^ we reasoned that they should be preferentially tuned to respond to sweet and fat relative to other gustatory cues. To test this, we recorded the single cell responses of DMH^LepR^ neurons in response to a panel of nine tastants that included bitter (quinine), sour (citric acid), salt (NaCl), fat (Intralipid), fat texture (silicone oil), and a complete diet (Ensure). We also included sucrose and sucralose as well as an alternative carbohydrate that has greatly reduced sweet taste (polycal).^47^ These solutions were presented in triplicate, in pseudo-randomized order while we recorded neural responses in food-deprived mice, and neurons were scored as responsive when they responded to a tastant in two out of three trials (Figure 4N).

Overall, we found that 89% of DMH^LepR^ neurons consistently responded to at least one tastant, and Ensure elicited responses in the largest percentage of neurons (59%). To characterize the diversity of responses, we used unsupervised k-means clustering to sort these neurons based on their activity across different solutions, which revealed five subpopulations with distinct response profiles (Figure 4OP). Two subpopulations had narrow response profiles: one that preferred sweet solutions, and one that preferred fatty solutions. Of note, the neurons that responded to Intralipid also responded to silicone oil, and similarly for sucrose and sucralose, indicating that these neurons respond not only to nutrients per se but also nutrient-associated tastes and textures. In addition, we observed three broadly tuned populations: one that had strong responses to a complete diet and weaker responses to pure tastants; a second that responded equally to all tastants; and a third, small cluster (n=3 neurons) with weak responses. Of note, there was no subpopulation that preferentially responded to salt, sour, or bitter, consistent with the idea that the function of taste in the DMH is to signal the presence of calories. This is achieved through dedicated subpopulations of LepR neurons that are tuned to respond to different combinations of nutritive tastes.

We noted that the subclusters above are biased towards specific tastes, but that their specificity is not absolute, similar to the mixed coding observed in the gustatory pathway.^48^ The metrics of noise-to-signal ratio and entropy are commonly used to quantify this specificity for tastants,^49,50^ and we applied this analysis to Ensure-responsive DMH^LepR^ neurons. Analysis of the noise-to-signal ratio, which measures the difference in response magnitude between the first- and second-best tastes, revealed that sweet-preferring neurons had the lowest noise-to-signal ratio, while the neurons preferring tastants not associated with calories (e.g. salt) had the highest ratio, indicating that sweet taste is encoded the most selectively (Figure 4Q). We also calculated the entropy for each of the neurons, which measures the breadth of tuning across many tastants. This revealed that the sweet- and fat-preferring neurons have the lowest entropy, meaning that they respond to the fewest number of tastants (Figure 4R). The relationship between these variables can be visualized by plotting the noise-to-signal ratio against entropy, which illustrates again that the most highly-tuned neurons in the DMH^LepR^ population are neurons that prefer sweet and fat taste (Figure 4S).

### DMH-LepR neurons integrate nutrient and gustatory signals

DMH^LepR^ neurons respond to the taste of food, but feeding behavior is also influenced by a food’s energy content, which is directly sensed in the GI tract.^51^ To probe this interaction between taste and calories, we characterized how DMH^LepR^ neurons respond to the same nutrient when it is either consumed by mouth or infused directly into the stomach.

We equipped mice with intragastic (IG) catheters for nutrient infusion into the stomach while simultaneously recording single-cell dynamics of DMH^LepR^ neurons (Figure 5A). We then infused Ensure (1.0 mL, approximating a moderately sized meal) into the stomach over ten minutes, which resulted in activation of 25% of DMH^LepR^ neurons (mean responses, 4.7 ± 1.1 z). This activation gradually ramped starting 49.7 ± 7.9 seconds after the start of infusion, reached a peak 4.5 ± 0.5 min later, and then slowly declined (tau = 20.0 ± 0.8 min). We observed similar sustained, ramping responses to IG infusion of sucrose or Intralipid (Figure S4). In contrast, IG infusion of a non-caloric, osmolarity-matched salt solution (1.0 mL over 10 min) activated fewer cells (4.1% vs 25%, p<0.0001; Figure 5D) and the magnitude of their activation was smaller (1.7 ± 0.3 z vs 4.7 ± 1.1 z, p = 0.041; Figure 5E), indicating that nutrients are required for the full response. Of note, we also detected some neurons that were inhibited by IG infusion (Figure 5B), but in this case there was no difference in the number of inhibited cells between saline and Ensure, (25% vs 29%, p=0.24; Figure 5D), and the difference in the magnitude of their inhibition was modest (-1.8 ± 0.1 z vs -1.4 ± 0.1 z, p=0.0002), indicating these inhibitory responses are mostly nutrient-independent.

**Figure 5.**
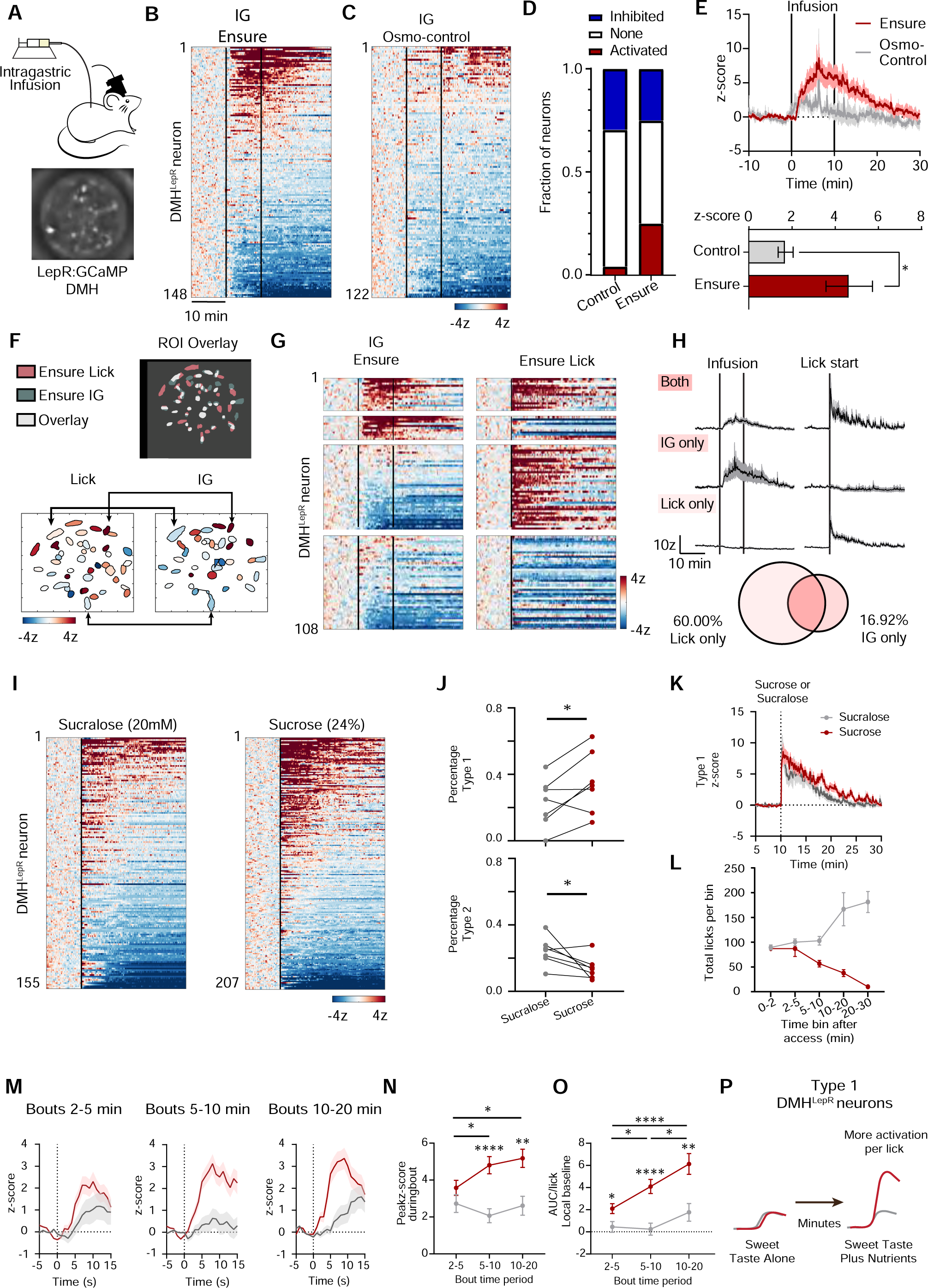
Nutrients potentiate DMH^LepR^ neuron responses to gustatory signals. **A,** Schematic and example of intragastric (IG) infusion setup during single-cell calcium imaging. **B,** Heatmap of DMH^LepR^ neurons (N=5 mice) receiving an IG infusion of ensure. **C,** Same as in B but with an osmolarity-matched control. **D,** Quantification of neural response types. **E,** Top, averaged trace of activated neurons. Bottom, mean z-score. **F,** Example overlay generated using CellReg to cross-register neurons. **G,** Heatmaps of aligned neurons, divided based on response patterns: activated by both, activated only by IG ensure or licking ensure, or other. **H,** Averaged traces from the categories in G, and corresponding percentage out of the number activated by at least one stimulus. **I,** Heatmaps during sucrose or sucralose consumption (N=7). **J,** Percentage of type 1 or 2 neurons per mouse. **K,** Averaged trace of type 1 neurons during consumption. **L**, Total licks per time bin. **M,** PSTH around licking bouts using a local baseline (N=4). **N,O,** Quantification of M without (N) and with (O) normalization to licks. **P,** Summary model of nutrients potentiating gustatory responses over time. Dots represent individual type 1 neurons. N.S. p>0.05, *p<0.05, **p<0.01, ****p<0.0001 Data is presented as mean ± SEM. See also Figure S4.

We next investigated the anatomic relationship between the DMH^LepR^ neurons that respond to IG versus oral Ensure. We did this by recording calcium dynamics while mice drank Ensure, or received a passive IG infusion, and then cross-registering cells between trials (Figure 5F). First, we noted that all cells activated during oral consumption showed short latency responses consistent with activation by licking, whereas the activation by IG infusion was invariably slower (time to peak 10.5 ± 0.4 vs. 269 ± 32s, p <0.0001), confirming that these are indeed two distinct responses. Second, we found that more neurons were activated by licking Ensure than IG infusion (50% vs 24%, p=0.0001) and, within this larger population of licking-activated neurons, 28% also responded to IG infusion. In contrast, within the smaller population of IG infusion-responsive cells, approximately half responded to licking (58%). Thus, there are subsets of DMH^LepR^ neurons that respond only to oral or gut signals, and a subset that responds to both.

Because many DMH^LepR^ neurons were activated by both licking and IG infusion, we examined how these signals are integrated at the single cell level (Figure 5F). The simple hypothesis that oral and GI signals are independent and additive was inconsistent with two observations. First, we found that a population of cells was activated by IG infusion of Ensure but not by oral ingestion (16.9% of all activated neurons). This is inconsistent with an additive mechanism, because all food consumed by mouth reaches the stomach. Second, there was no difference in the magnitude or duration of the activation DMH^LepR^ neurons during oral ingestion that correlated with whether the cells also responded to IG infusion (Figure 5G). For example, there was no significant difference in the mean activity of neurons activated by licking only or by both licking and IG Ensure (5.6 ± 0.97 vs. 6.7 ± 1.9z, p=0.66). This is inconsistent with the prediction of an additive mechanism, because cells responding to both signals should be activated more strongly.

We next considered the alternative hypothesis that, during normal ingestion, nutrient signals function to modulate the gain of responses to orosensory cues. To test this, we measured how lick responses to sucrose and sucralose evolve over longer timescales (30 min), and in the presence of higher levels of consumption, in order to expose a possible role for post-ingestive feedback. Of note, we found in Figure 4B,C that DMH^LepR^ neurons are activated similarly by sucrose and sucralose ingestion in brief-access tests of 5 second duration (2.8 ± 0.4 z vs. 2.5 ± 0.5 z, p=0.9), but those experiments, by design, do not allow for post-ingestive feedback.

Mice were given access to either sucrose or sucralose for 30 minutes and we examined the response of DMH^LepR^ neurons to self-paced consumption (Figure 5I). We found that consumption of sucrose and sucralose resulted in broadly similar patterns of DMH^LepR^ neurons activation over the 30-minute session (Figure 5K), with two important differences. First, drinking sucrose was associated with a higher percentage of Type 1 neurons (cells activated by licking throughout the trial; Figure 5J) and a smaller percentage of Type 2 neurons (cells activated only at the trial start; Figure 5J) compared to drinking sucralose. This suggests that caloric solutions recruit more cells that are durably activated by ingestion. Consistent with this, we noticed that the neural response to licking sucrose became progressively stronger as the trial progressed, whereas the response to sucralose was unchanged. This can be visualized by plotting the mean response during lick bouts at different stages in the trial (2-5 min, 5-10 min, and 10-20 min; Figure 5M) and can be quantified by measuring the peak z-score during these bouts, which was initially the same but then diverged as the trial progressed (mean diff. -0.85 (2-5 min), -2.7 (5-10 min), and -2.6 (10-20 min), Figure 5N). Thus, the time-locked activation during licking sucrose, but not sucralose, increases over the course of a meal, and this effect emerges on a timescale (5-20 min) consistent with a role for post-ingestive nutrient feedback in amplifying responses to orosensory cues.

An alternative explanation for these findings is that they are secondary to changes in behavior: if the mice lick more for sucrose as the trial progresses, then DMH^LepR^ neurons may appear to be more activated by sucrose for this reason alone. However, when we analyzed the licking data, we found that the opposite was true: as the 30-minute session progressed, sucrose intake decreased (presumably due to satiety), while sucralose intake increased ((F_Time X Solution_(1.417,4.25) = 14.25, p=0.015; Figure 5L). As a result, when we normalized neural responses to the number of licks in the bout, the divergence between sucrose and sucralose was even greater (F_Solution_ (1,85) = 15.75, p=0.0002; Figure 5O). Taken together, these results suggest that post-ingestive feedback modulates DMH^LepR^ neurons, at least in part, by potentiating their lick-evoked activation by sweet tastes (Figure 5P). This provides a mechanism by which orosensory and gastrointestinal cues could interact to synergistically inhibit appetite.

## Discussion

The sense of taste provides the first detailed assessment of the quantity and quality of ingested food. In contrast to nutrient sensing in the small intestine, which is inherently delayed by slow gastric emptying,^52^ gustatory feedback is immediate, enabling a forecast of the nutritional effects of ongoing ingestion. Thus, it would have obvious adaptive value for the nervous system to use these gustatory cues, not only for food discrimination^53–55^ and reward,^35,56,57^ but also to initiate the satiation processes that will ultimately lead to meal termination.^58^

Consistent with this logic, there have been behavioral observations suggesting a role for orosensory cues in promoting satiation. These include the fact that food is often more satiating when consumed by mouth than when delivered directly to the gut;^3–7^ that loss of GI feedback during sham feeding does not eliminate all of the satiating effects of food ingestion;^10,12,59,60^ and that repeated exposure to specific tastes can reduce their consumption, independent of any GI cues.^13,14,61^ Still, the idea that taste functions as an important satiation signal has achieved little traction in the scientific community. A fundamental obstacle has been the difficulty of separating the contribution of food tastes to the termination of feeding from their critical and inescapable role in promoting the initiation and maintenance of ingestion.

Here, we have taken a neural dynamics approach to disentangling these opposing functions of taste cues on behavior. Our strategy has been to examine how food tastes modulate neurons that are genetically hard-wired to promote feeding, and then ask what happens when this gustatory modulation is blocked. We have shown that AgRP neurons, the most widely studied cells that promote hunger, are inhibited by food tastes every time a mouse takes a lick of liquid diet. Moreover, we found that the taste of food is the dominant signal that regulates the direct GABAergic afferents to AgRP neurons, DMH^LepR^ neurons, during a normal meal. For both of these cell types, the sign of their modulation implies a role for taste cues in inhibiting food intake, and, consistently, we found that blocking this gustatory modulation increases food intake by specifically delaying satiation. These findings establish a neural substrate for the negative feedback control of ingestion by the taste of food, opening the door to systematic study of this fundamental but elusive phenomenon.

### AgRP neurons track and modulate the dynamics of ingestion

Given their central role in hunger, there is considerable interest in understanding how AgRP neurons are regulated. Early studies showed that AgRP neurons are rapidly inhibited at the start of a meal by the sight and smell of food.^17,20,24^ The magnitude of this inhibition predicts the amount of food subsequently consumed, suggesting that the primary regulation of AgRP neurons during feeding is anticipatory in nature and occurs before the meal begins.^25,27^ Nevertheless, AgRP neurons do show ongoing activity during ingestion, albeit at a reduced level, and it has remained an open question what drives these intrameal dynamics^17,24^ and whether they have any significance for behavior.

To address this, we reinvestigated the dynamics of AgRP neurons using new calcium sensors that are more sensitive to activity fluctuations at low firing rates,^62^ and therefore more likely to detect fluctuations in AgRP neuron activity during the time period immediately after food discovery, when overall activity is low. This revealed a dramatic, time-locked inhibition of AgRP neurons during each feeding bout that is triggered by contact with food and strongest for food-associated tastes such as sweet and fat. In contrast to the AgRP neuron response to GI feedback,^25,63^ this lick-evoked inhibition does not require calories (Fig. 1), and in contrast to the rapid inhibition by sight and smell of food,^17,20,24^ it does not undergo extinction in the absence of nutrients (Fig. S1). This reveals that the taste of food plays a unique role in driving AgRP neuron activity in the time interval between initial food discovery and later post-ingestive feedback.

Optogenetic reversal of the taste-specific modulation of AgRP neurons resulted in an increase in the number of feeding bouts, with no effect on their size. This increase was specifically manifest as a slowing of the decline in the rate of bout initiation that naturally occurs as a meal progresses and animals approach satiety. This suggests that the ingestion-triggered dips in AgRP neuron activity are sensed by downstream circuits in order to modulate the timing of meal termination. We do not know which downstream circuits are the target of this signal or how their activity is modulated, but the paraventricular hypothalamus, bed nucleus of the stria terminalis, lateral hypothalamus, and paraventricular thalamus are known to be important for AgRP neurons’ effects on feeding.^24,64,65^ It is also unknown whether this gustatory inhibition of AgRP neurons exclusively modulates caloric satiety or if it also plays a role in sensory-specific satiety to individual tastes. These are important questions for future investigation.

In contrast to the closed-loop stimulation described here, previous studies have shown that tonic, high-frequency stimulation of AgRP neurons can drive increases in appetite that persist for at least an hour, even in the absence of continued stimulation.^21^ This long-lasting response to AgRP neuron “prestimulation” is mediated by NPY^32^ and is distinct in several ways from the response to closed-loop stimulation described here, which (1) requires precisely-timed light delivery, (2) has effects that extinguish quickly with the offset of laser stimulation, and (3) is achieved using low-frequency stimulation that is inefficient inducing neuropeptide release.^66–68^ Moreover, prestimulation of AgRP neurons increases bout size with no effect on bout number,^32^ whereas closed-loop stimulation described here does the opposite. Consistent with its effect on bout size, which tracks palatability, prestimulation of AgRP neurons conditions robust flavor preference,^21^ whereas the closed-loop stimulation here does not (Fig. 2). This reveals that AgRP neuron activity before and during feeding control fundamentally different aspects of ingestion.

### DMH^LepR^ neurons encode a representation of appetitive tastes

The DMH is a structure traditionally associated with the regulation of autonomic responses (cardiac output, energy expenditure, body temperature) and behavioral rhythms (circadian and food entrainment).^69–73^ In addition, DMH^LepR^ neurons have been implicated in the control of food consumption and learning about food cues via their direct projection to AgRP neurons.^26,44^ Population recordings using fiber photometry have reported that these DMH^LepR^ neurons can be activated by the sight and smell of food, suggesting that they drive the “sensory drop” in AgRP neuron activity that occurs when a meal begins. However, the finding that AgRP neurons are inhibited by food tastes during ingestion motivated us to reexamine the dynamics of these cells using single-cell imaging.

This revealed that the DMH^LepR^ neuronal population contains three subsets of neurons that are modulated during feeding in distinct ways. One of these subsets is rapidly and transiently activated when food is first presented, and therefore could contribute to the rapid inhibition of AgRP neurons when a meal begins.^17,20,24,26,44^ However, we found that the largest population of modulated cells, and the only population that responded specifically to food, was transiently activated during licking in a manner time-locked to ingestion. These responses scaled with the intensity of food-associated tastes, and at the single-cell level distinct neurons were tuned to either sweet or fat, while others responded more broadly. Of note, tracing studies have shown that the DMH receives input from regions known to respond to taste cues,^44,74,75^ and a gustatory input to these circuits is also supported by the results of polysynaptic retrograde tracing from AgRP neurons, which revealed prominent input from classic taste regions such as the rostral nucleus of the solitary tract and parabrachial nucleus.^76^ Thus, taste is an important source of sensory feedback to the DMH and may be involved in regulating DMH outputs beyond feeding behavior.

## Acknowledgements

This work is supported by NIH grants R01DK106399 (Z.A.K.), RF1NS116626 (Z.A.K.), F31DK125067 (T.J.A.) and University of California, San Francisco William K. Bowes, Jr. and Ute Bowes Discovery Fellowship (T.J.A.). Z.A.K. is an HHMI Investigator.

## Author Contributions

Conceptualization, TJA, and ZAK; Methodology: TJA, TL, NLSM, LAG, and ND; Investigation: TJA, SS, NS, and ZAK; Validation: TJA and CB; Visualization: TJA and ZAK; Funding Acquisition: TJA and ZAK; Project administration: TJA and ZAK; Supervision: ZAK; Writing—original draft: TJA and ZAK; Writing—review & editing: TJA, TL, and ZAK.

## STAR Methods

### Resource availability

#### Lead contact

Further information and requests for resources and reagents should be directed to and will be fulfilled by the lead contact, Zachary A. Knight.

#### Materials availability

This study did not generate any new unique reagents.

#### Data and code availability

All data reported in this paper will be shared by the lead contact upon request.

All original code has been deposited on GitHub and is publicly available as of the date of publication.

Any additional information required to reanalyze the data reported in this paper is available from the lead contact upon request.

### Experimental model and subject details

#### Animals

LepR-IRES-Cre (Jackson #032457, RRID: IMSR_JAX:032457), AgRP-IRES-Cre (Jackson #012899, RRID: IMSR_JAX:012899), and AgRP-IRES-Cre crossed with Ai32:ROSA26-loxSTOPlox-ChR2-eYFP (Jackson #012569, RRID: IMSR_JAX:012569) adult (>6 weeks) mice of both sexes were used for experiments. Mice were kept in a humidity and temperature-controlled housing facility on a 12-hour light/dark cycle and had ad libitum access to food (PicoLab 5053) and water unless otherwise noted for experiments. All LepR-Cre experimental mice were singly housed for experiments. For fasted and water-deprived experiments, mice were food-deprived or water-deprived overnight before the experiment, respectively. All mice were habituated to the experimental chamber overnight, and mice were habituated to being handled at least one day before experiments. Littermate Cre-negative, Rosa-negative, or wildtype (Jackson #000664, RRID: IMSR_JAX:000664) controls were used where possible. All behavioral protocols were approved by University of California, San Francisco’s Institutional Animal Care and Use Committee.

## Method details

### Stereotaxic Surgeries

For all stereotaxic surgeries, mice underwent procedures as described previously ^24^. Briefly, mice were anesthetized using isoflurane. Following surgery, mice were left for 1-4 weeks for recovery and viral expression.

For microendoscopic imaging experiments, LepR-Cre mice were stereotaxically injected with 150-200 nL of AAV1-CAG-Flex-GCaMP6s-WPRE-SV40 (6.1 × 1012 titer; Addgene) unilaterally into the left DMH (AP: -1.8 ML: -0.4 DV: -5.2 or -5.3), and a GRIN lens (8.4 × 0.5mm; Inscopix 1050-004610) was implanted 0.05 mm medial and 0.1 mm dorsal to the injection site. The lens was secured to the skull with metabond dental cement (Parkwell S380). Following 4 weeks to allow for virus expression, mice were anesthetized again, and a baseplate (Inscopix 100-000279) was affixed above the lens with metabond and covered with a baseplate cover (Inscopix 100-000241).

For fiber photometry, AgRP-Cre mice were stereotaxically injected with AAV1-CAG-FLEX-GCaMP8s (1 × 1013 titer; Janelia) into the ARC (AP: -1.8 ML: -0.35 DV: -5.8), and an optical fiber (inner diameter 0.4 mm by 8 mm in length; Doric lenses MFC_400/430-0.48) was placed 0.05 mm medial and 0.1 mm dorsal to the injection site.

For optogenetic experiments, AgRP-Cre::Rosa-ChR2 and control mice (C57bl/6 wildtype, AgRP-Cre, or Rosa-ChR2) were stereotaxically installed with custom-made fiber optic implants (0.39 NA Ø200 µm core Thorlabs FT200UMT and CFLC230-10) unilaterally above the ARC (AP: -1.8 ML: -0.3 DV: -5.7-5.8).

### Intragastric Catheterization

Mice were equipped with an intragastric catheter as described previously ^25,77^. Briefly, mice were anesthetized with ketamine-xylazine and a sterile veterinary gastric catheter (C30PU-RGA1439, Instech Labs) was surgically implanted into the avascular forestomach through the abdominal wall. The catheter was attached to a sterile access button (VABM1B/22, Instech Labs), which was implanted between the shoulder blades of the mouse’s back. The port was protected with an aluminum cap (VABM1C, Instech Labs) that was placed between experiments. Mice were allowed one week to recover before performing experiments.

Catheters were flushed the night before a recording session with deionized water to ensure patency. Intragastric infusions consisted of an infusion of vanilla Ensure (1 mL, 0.32 g/mL) (Abbott) or 225 mM NaCl over 10 minutes at a rate of 100 μL/min.

### Microendoscopic Imaging

All data was recorded using the Inscopix nVista (v. 3.0) or nVoke (v2.0) systems. Data was acquired using the Inscopix acquisition software (v151) at 20Hz, 8 gain, 0.5-0.7 mm^−2^ 455 nm LED power, and 1-2x spatial downsampling. Prior to all recordings, mice were attached to the cameras in the chambers with the LED on for 10 minutes to allow for habituation and stabilization of GCaMP signal. Mice presented with peanut butter were given continuous access for 30 minutes following the conclusion of the experiment. Consumption of liquid food and fluids was monitored throughout the 30-minute experiment using a contact lickometer constructed in-house. Individual bites of peanut butter were hand-scored from video footage taken during the experiment.

### Fiber photometry

Fiber photometry experiments were conducted as described previously ^78^. The fiberoptic implant was cleaned with 70% alcohol and a cleaning stick (MCC25) before mice were tethered to a patch cable (Doric Lenses, MFP_400/460/900-0.48_2m_FCM-MF2.5), and were given 10 minutes to habituate prior to the experiment. A 6-mW blue LED (470 nm) and UV LED (405 nm) served as continuous light sources throughout. These light sources were driven by a multichannel hub (Thorlabs), modulated at 305 Hz and 505 Hz, respectively, and delivered to a filtered minicube (Doric Lenses, FMC6_AE(400–410)_E1(450–490)_F1(500–540)_E2(550– 580)_ F2(600–680)_S) before transmitting through a patchcord and implanted optic fibers. GCaMP and isosbestic UV fluorescent signals were collected through the same fibers back to a minicube and into a femtowatt silicon photoreceiver (Newport, 2151). Digital signals were sampled at 1.0173 kHz, demodulated, lock-in amplified, sent through a processor (RZ5P, Tucker-Davis Technologies (TDT)), and collected by the provided software Synapse (TDT). Data was lastly exported through Browser (TDT) for following analysis in MATLAB.

Mice were presented with different solutions connected to a custom-made contact lickometer for thirty minutes following an additional 10-minute baseline period. For injections, mice were first injected with vehicle (saline) or devazepide (1 mg/kg, R&D Systems) i.p. and intralipid access was given five minutes later. Mice were excluded if they did not have a sensory drop in AgRP activity at food access.

### Davis Rig

A Davis Rig gustometer (MED-DAV-160M, Med Associates) was used to perform brief access taste tests in mice. This rig is equipped to hold 16 bottles on a motorized plate that can position individual bottles in front of an entry port. The entry port is closed off using an automated door. Mice were trained over two days while water deprived overnight. The first day, mice were allowed to freely lick from one waterspout for thirty minutes. The second day, mice were allowed intermittent access to a waterspout. Mice were given ad libitum access to water for at least 2 hours between water deprivations. Subsequent testing was done in overnight food-deprived mice.

Sucralose and sucrose taste curves were performed using 1:2 serial dilutions from 0.32 g/mL (32%) sucrose and 25mM sucralose solutions. This totaled 9 bottles with the sweetener, and 3 bottles of water were added as a control. The 12 bottles were presented in pseudo-random order 3 times, yielding 36 total trials.

Taste panel tests consisted of 12 total bottles, where 3 bottles contained water and 1 bottle each of: 0.32 g/mL Ensure, 8% polycal (Nutricia), 20% Intralipid (Sigma Aldrich I141-100mL, Medline BHL2B6064H), 12.5 mM Sucralose, 100 mM NaCl (salt), 20mM citric acid, silicone oil (Sigma Aldrich 378348), 16% sucrose, and 0.3 mM quinine. The 12 bottles were presented in pseudo-random order 3 times, yielding 36 total trials. The first lick in a presentation was synced to the nVista system via a single TTL pulse. Licking activity was recorded and outputted as a separate file from the Med Associates program.

### Optogenetics

Mice were functionally validated by stimulating (20 Hz, 15 mW, 10 ms pulse width, 2 s on/3 s off) fed mice for one hour during access to chow. Mice were included in the experiment if they ate at least 0.6 g of chow.

The Coulbourn Habitest system was used to control the 473 nm laser in a setup described previously ^21^. Paired and unpaired 2-minute trials were randomly assigned with 50% probability for each throughout the 1-hour session. During paired trials, licks detected by a custom-made contact lickometer triggered 2-s of stimulation (5 Hz, 15 mW, 10 ms pulse width). Laser stimulation always ended 2-s after the last lick, such that overlapping licks did not cause extended stimulation times. No laser was given during unpaired trials, but licks were recorded.

Conditioned flavor preference was conducted using the same Coulbourn Habitest system. Mice were tethered throughout training and testing. Mice were first habituated to the individual test solutions (Lime or Grape 0.05% Koolaid with 16% Sucrose) over two days. A baseline two-bottle preference test was then done over two days, where both bottles were presented, their order switching the next day. The next four training days consisted of alternating pairing one solution with closed-loop stimulation (2-s per lick, 5Hz, 15 mW, 10 ms pulse width) and the other solution paired with nothing. The two-bottle preference test was then repeated over two days. Test order and flavor pairings were counterbalanced across mice.

### Histology

Mice were anesthetized and transcardially perfused with phosphate buffered saline (PBS) and 10% formalin, as described previously ^78^. Brains were stored in 10% formalin overnight at 4°C and moved to 30% sucrose in PBS the next day. Two days later, brains were sectioned (40µm) using a cryostat and mounted onto slides with DAPI Fluoromount-G (Southern Biotech). Viral expression and lens placement was then evaluated using a confocal microscope.

### Quantification and statistical analysis

#### Data Analysis

All data analysis was conducted in MATLAB unless otherwise noted.

Fiber photometry data was normalized using the function: ΔF/F_0_ = (F-F_0_)/F_0_ as described previously ^78^, where F is the raw photometry signal and F_0_ is the predicted fluorescence using the 405 nm signal. Calcium traces and lick TTLs were finally downsampled to 1-Hz for presentation clarity. PSTHs were calculated for bouts at least 4-s long and 2-s apart, excluding data from the first two minutes to remove the contribution of the sensory drop in AgRP activity. Point-biserial correlation coefficients were calculated by transforming the lick TTLs into a binary trace and correlating this with the analyzed calcium trace using R. Again, lick and calcium activity during the first two minutes of presentation was excluded to remove the contribution of the sensory drop in activity. Time constants were calculated by finding the time for the signal to decay to 37% of the peak. Means for activity across the whole trial were taken during the first 10 minutes after presentation.

Optogenetic behavior data was analyzed using custom-written code. Bouts were defined as being at least 2-s long and at least 2-s apart. Bouts that extended across trials were excluded. Preference for a flavor was calculated by dividing the two-day average licks for the paired flavor by the two-day average total licks.

Single-cell calcium imaging data was first preprocessed using the Inscopix Data Processing software (v1.3.1) to spatially (binning factor of 2 or 4) and temporally (binning factor of 2) downsample the videos. These were then put through a spatial bandpass filter to remove noise and out-of-focus cells. Finally, videos were motion-corrected using the above software or Inscopix Mosaic software (v1.2, to correct non-translational motion artifacts). Cells or whole videos were removed if the motion could not be corrected. Videos were then sent through the constrained non-negative matrix factorization for endoscopic imaging (CNMFe; ^79^) pipeline to extract activity traces of single neurons. ROIs were removed if they were duplicates, their activity patterns were influenced by surrounding fluorescence, encompassed the edge of the lens, or were an over-segmentation of a larger ROI.

For experiments with free access to a solution or an intragastric infusion, traces were z-scored to the 10-minute baseline period prior to access. For ensure and chow consumption experiments, DMH^LepR^ neurons were separated into 4 categories based on their activity patterns. “Type 1” or “Ingestion” was defined as having a mean over the first 10-minutes greater than or equal to 1. Similarly, “Type 3” or “Inhibition” was defined as having a mean over the same time scale less than or equal to -1. “Type 2” or “Presentation” neurons had an average z-score in the first 60-seconds greater than or equal to 1, but less than 1 over the first 10-minutes. PSTHs were generated by cutting activity 15-seconds pre and post the start of a bout (at least 4 seconds of licking, with at least 2 seconds between bouts), and z-scoring activity to the pre-bout baseline. Onset latency was calculated by finding the time post first lick when activity crosses a threshold of 1z. Correlations were made by turning the lick timestamps into a binary “trace” and computing the point-biserial correlation coefficient for each neuron in the respective category.

For davis rig experiments, single-cell calcium traces were processed through the CNMFe pipeline described above. Calcium traces were cut around the 15 seconds before and after the first lick in a presentation. Neurons were deemed activated by sucrose or sucralose if the mean activity after the first lick is greater than or equal to 1 for at least two of the three trials of second-to-maximal concentration. Similarly, for the taste panel, neurons activated by ensure the majority of the time were categorized into different response types using unsupervised k-means in MATLAB. Entropy was analyzed using neural responses to a subset of basic tastants: sucralose, silicone oil, salt, citric acid, quinine, and water. Entropy was calculated using the formula 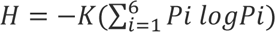 where K = 1.2851 for 6 tastants ^49^. Noise to signal ratio was calculated by dividing the second maximal response by the maximal response ^50^. Activity traces across free licking and intragastric infusions of ensure were cross registered across days using previously published CellReg MATLAB code ^80^.

#### Statistics

Statistical values (e.g., N mice, n neurons, p, F) can be found in the results section, figures, and figure legends. Values are reported as mean ± sem (error bars or shaded areas. *P*-values for comparisons across multiple groups were calculated using one- or two-way ANOVAs, where appropriate, and following multiple comparisons were calculated using Tukey or Sidak tests, respectively. One-sample comparisons to a null hypothesis value of 0 were done using a one-sample two-tailed t test. Direct comparisons between two samples were tested using either a t-test, Mann-Whitney, or Wilcoxon test, where appropriate. Comparisons of model fits was determined using an Extra sum-of-squares F test, where a linear model was compared to a quadratic model (Figure 4H,M). Percentage comparisons in Figure 5D and G were calculated using a Fisher’s exact test. All statistical tests were performed in GraphPad Prism 8, except for a K-S test performed in MATLAB and point-biserial correlations performed in R. Power calculations were used to determine sample size when possible, based on preliminary data availability (Figures 2, 3 and 5). Experiments were not randomized, but bottle position and paired flavor were counterbalanced for conditioned flavor preference experiments. Blinding was not used while conducting experiments or performing analyses.

## Supplemental information titles and legends

**Supplementary Figure 1 – related to Figure 1.**
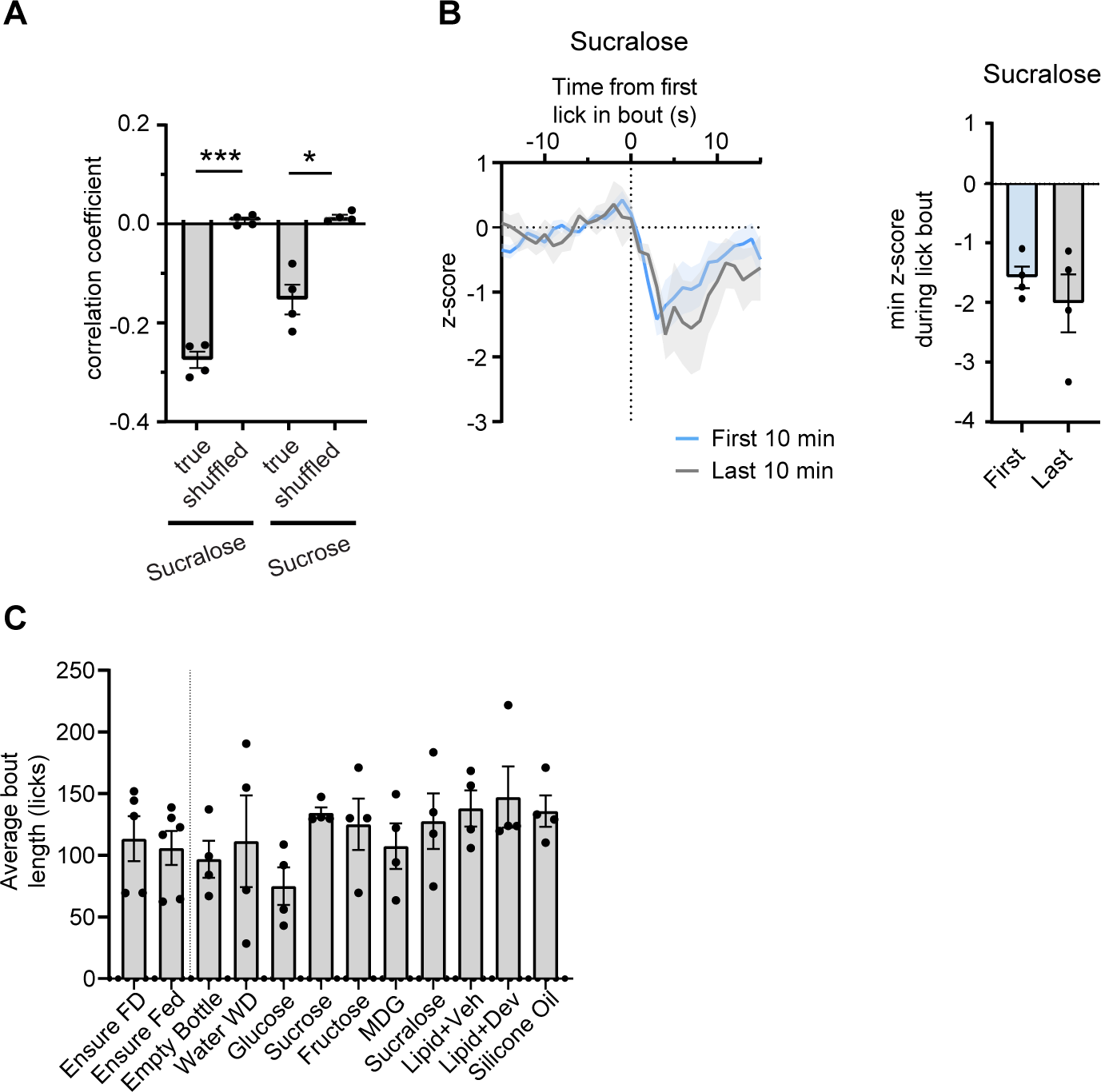
AgRP lick-evoked activity during sucralose consumption does not decay over time. **A**, Point-biserial correlation coefficient when comparing AgRP calcium activity with either the true or shuffled lick activity for sucralose and sucrose. **B**, PSTH of lick-evoked AgRP activity when drinking sucralose during either the first or last 10 minutes of a 30-minute session (left), and corresponding quantification (right). **C**, Average bout lengths during PSTHs shown in Figure 1. N.S. p>0.05, *p<0.05, ***p<0.001. Data is presented as mean ± SEM.

**Supplementary Figure 2 – related to Figure 3.**
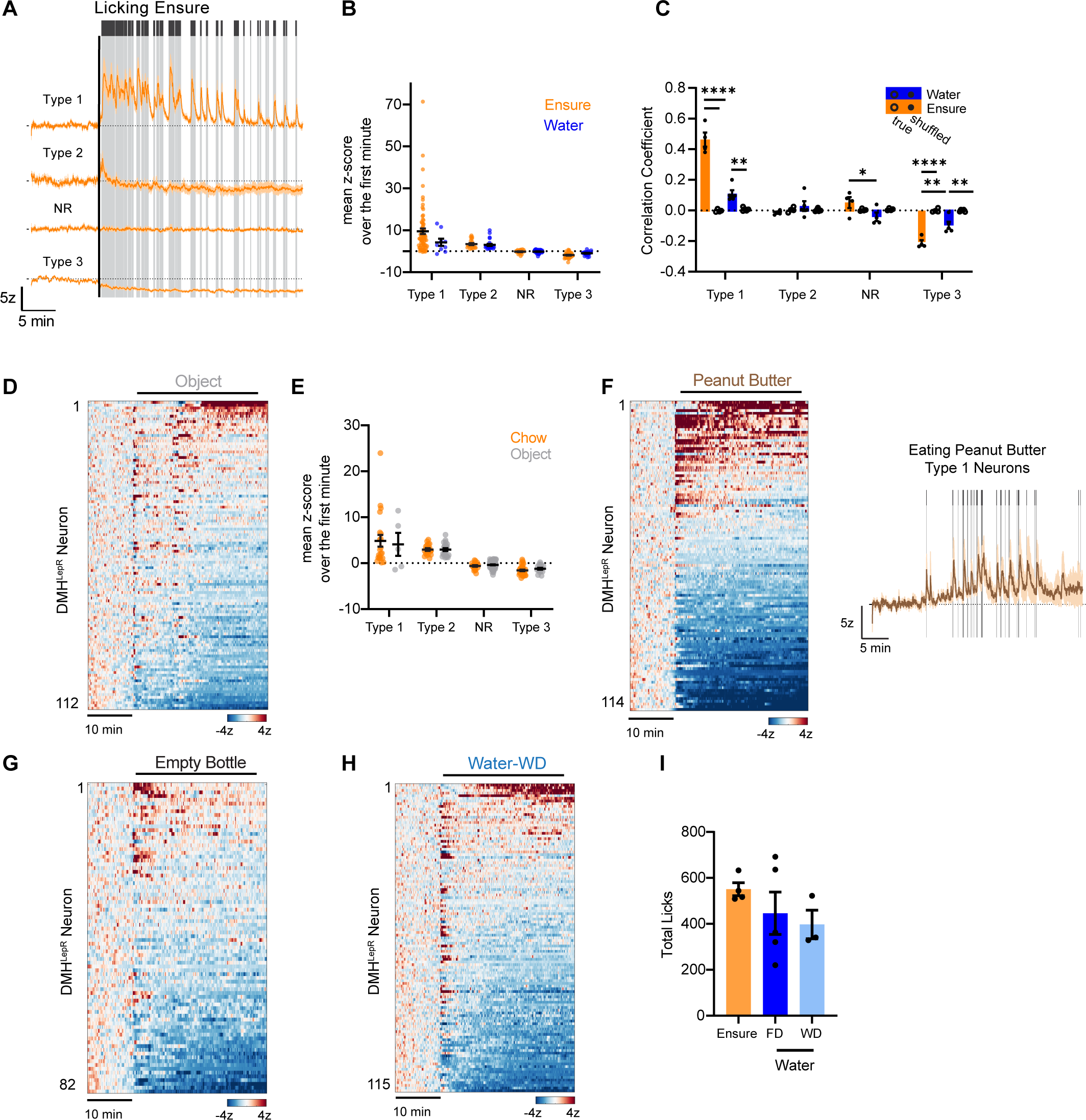
DMH^LepR^ activity during consumption. **A**, Example averaged trace of all 4 types of responses during licking ensure. **B**, Mean z-score over the first minute for each response category. Dots represent individual neurons. **C**, Correlation coefficient for neurons across all response categories comparing their responses to either true or shuffled licking activity. **D**, Heatmap of neurons from 3 mice given an inedible object. **E**, Mean z-score over the first minute for neurons across all response categories during chow ingestion or object presentation. **F**, Heatmap of neurons from 4 mice eating peanut butter. Example averaged activity of type 1 neurons shown right, with bites marked in grey. **G**, Heatmap of neurons from 3 mice licking an empty bottle. **H**, Heatmap of neurons from 3 mice licking a water bottle while water deprived. **I**, Total licks for sessions of licking ensure or water. FD = food deprived, WD = water deprived. N.S. p>0.05, *p<0.05, **p<0.01, ****p<0.0001 Dots represent individual mice unless otherwise noted. Data is presented as mean ± SEM.

**Supplementary Figure 3 – related to Figure 3.**
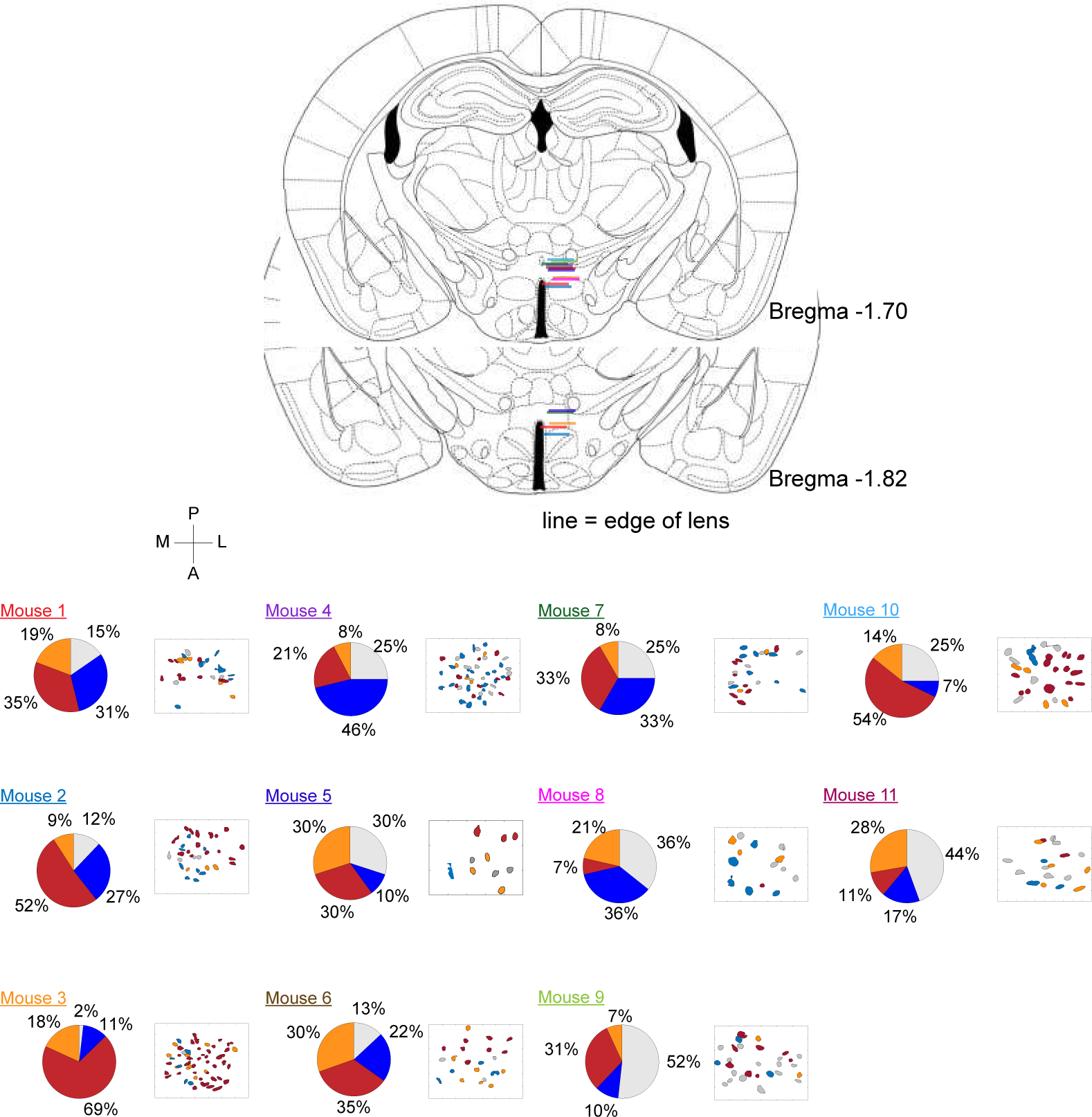
Schematic of lens placements and activity patterns across mice. Top, schematic representation of lens placement (colored line) for every mouse used in experiments shown in figures 3-5. Each color represents a different mouse. Bottom, Field of view schematic with neurons colored according to response type during ensure (mouse 1-4, 6), peanut butter (mouse 5), or sucrose (mouse 7-11) consumption. The color of the mouse label corresponds to the color of the lens placement bar above. The pie chart represents proportions of neurons for each mouse fitting into each category. Red = type 1, orange = type 2, grey = nr, blue = type 3.

**Supplementary Figure 4 – related to Figure 5.**
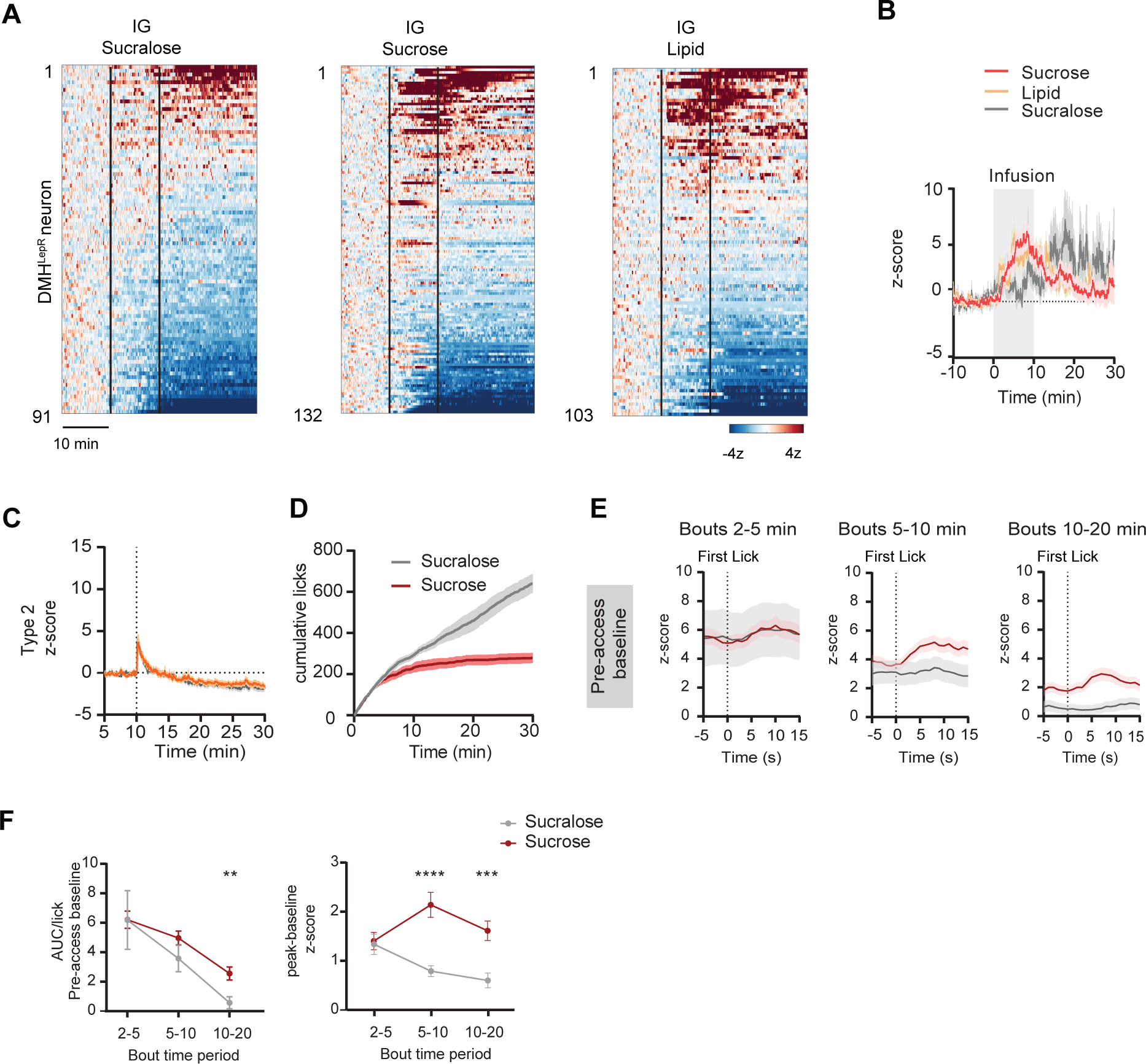
DMH^LepR^ responses to nutritive solutions. **A**, Heatmaps of DMH^LepR^ neurons during an intragastric (IG) infusion of sucralose (N=5 mice), sucrose (N=6), or intralipid (N=4). **B**, Mean traces of activated neurons during the infusions listed in a. **C**, Mean trace of type 2 neurons during sucrose (orange) or sucralose (grey). **D**, CDF of licks throughout consumption of sucrose or sucralose. **E**, PSTH of neural activity peri bouts in Figure 5M using a pre-access baseline. **F**, Quantification of activity depicted in E normalized to licking activity (left) and looking at the change during licking (right). N.S. p>0.05, **p<0.01, ***p<0.001, ****p<0.0001 Data is presented as mean ± SEM.

